# An electrophysiological approach to measure changes in the membrane surface potential in real time

**DOI:** 10.1101/607945

**Authors:** Verena Burtscher, Matej Hotka, Michael Freissmuth, Walter Sandtner

## Abstract

Biological membranes carry fixed charges at their surfaces. These arise primarily from phospholipid head groups. In addition, membrane proteins contribute to the surface potential with their charged residues. Membrane lipids are asymmetrically distributed. Because of this asymmetry the net negative charge at the inner leaflet exceeds that at the outer leaflet. Changes in surface potential are predicted to shape the capacitive properties of the membrane (i.e. the ability of the membrane to store electrical charges). Here, we show that it is possible to detect changes in surface potential by an electrophysiological approach: the analysis of cellular currents relies on assuming that the electrical properties of a cell are faithfully described by a three-element circuit - i.e. the minimal equivalent circuit - comprised of two resistors and one capacitor. However, to account for changes in surface potential it is necessary to add a battery to this circuit connected in series with the capacitor. This extended circuit model predicts that the current response to a square-wave voltage pulse harbors information, which allows for separating the changes in surface potential from a true capacitance change. We interrogated our model by investigating changes in capacitance induced by ligand binding to the serotonin transporter (SERT) and to the glycine transporters (GlyT1 and GlyT2). The experimental observations were consistent with the predictions of the extended circuit. We conclude that ligand-induced changes in surface potential (reflecting the binding event) and in true membrane capacitance (reflecting the concomitant conformational change) can be detected in real time even in instances where they occur simultaneously.

**Statement of Significance:** The plasma membrane of a cell possesses fixed charges on both surfaces. Surface charges play an important role in many biological processes. However, the mechanisms, which regulate the surface charge densities at the plasma membrane, are poorly understood. This is in part due to lack of experimental approaches that allow for detecting changes in surface charges in real time. Here, we show that it is possible to track alterations in the electric potential at the membrane surface with high temporal resolution by an electrophysiological approach. Importantly, the described method allows for discriminating between a change in surface potential and a change in true membrane capacitance (e.g. a change in membrane area), even if these occur in parallel.

## Introduction

Membrane capacitance recordings are primarily suited for measuring the plasma membrane surface area and changes thereof, with exquisite sensitivity (1, 2). For instance, techniques to measure the capacitance are ideally suited for this purpose: it is possible to detect a change in surface area resulting from the fusion of a single vesicle with the plasma membrane (3). However, it has long been known that membrane proteins, can also contribute to the measured membrane capacitance (4, 5). For instance, in membranes containing a large number of voltage-gated ion channels, the measured capacitance is comprised of two components: a voltage-independent component that can be ascribed to the membrane and a voltage-dependent component, which arises from mobile charges present in the voltage-sensing domains of the channels. Similarly, capacitance measurements also allow for monitoring conformational rearrangements that occur in membrane transporters. It has, for instance been possible to infer the conformational transition associated with phosphorylation of the Na^+^/K^+^ -pump or to track the conformational rearrangement associated with Cl^−^ binding to the inward facing conformation of the γ-aminobutyric acid transporter-1 (GAT1/SLC6A1)(6).

In all of the above cases, the observed changes were assumed to reflect true changes in the membrane capacitance arising from alterations in the polarizability of membrane embedded proteins. Macroscopically, these alterations can be described as a result of changes in relative permittivity of the protein (i.e. dielectric constant). Theoretical estimates for the relative permittivity of folded proteins fall into the range between 2.5 and 4 (7). Different conformations of a protein may exhibit differences in their polarizability: if this is the case, the conformational rearrangement is expected to be measurable by capacitance recordings. However, alterations in the permittivity of a protein are not the only cause for a protein-associated capacitance change. We previously reported that the application of cocaine to HEK293 cells stably expressing the human serotonin transporter (hSERT) resulted in a reduction of membrane capacitance. We hypothesized that the change evoked by cocaine originated from a change in the outer surface potential caused by binding of the charged ligand to SERT rather than from a true capacitance change. Accordingly, we referred to this alteration as an apparent change in capacitance (8).

Cellular currents are usually assumed to be produced by a three-element circuit comprised of a capacitor (C_M_) representing the membrane and two resistors (R_S_ and R_M_) representing the series resistance and membrane resistance, respectively (i.e. minimal equivalent circuit of a cell) (9). However, the three-element circuit cannot account for the apparent change in capacitance that occurs as a consequence of charged ligand binding to a membrane protein. Here we show that cellular currents are better described by an extended circuit, which contains an additional battery. This battery, which is connected in series with the capacitor, can account for the ligand-induced change in surface potential. This allows for modeling the apparent capacitance change. Importantly, the extended circuit also predicts differences in the currents produced by a true and an apparent capacitance change. We verified these predictions by documenting that ligand-induced alterations and true capacitance changes can be differentiated based on information contained in current responses to a square-wave voltage stimulus. Importantly, our findings show that the measured current responses reveal information that allows for monitoring changes in the surface potential of the cell and in membrane capacitance also in those instances, where they occur simultaneously.

## Methods

### Cell culture

Human serotonin transporter (hSERT) tagged with GFP was stably expressed in a tetracycline-inducible HEK293 cell line. HEK293 cells have been previously authenticated by STR profiling at the Medical University of Graz (Cell Culture Core Facility). *Cercopithecus aethiops* SV40-transformed kidney (Cos-7) cells were obtained from ATCC (ATCC® CRL-1651). Cos-7 were transfected with a plasmid encoding N-terminally GFP-tagged hGlyT1b or hGlyT2a, respectively; the cDNAs of hGlyT1b in eGFP-C-1 and hGlyT2b in pcDNA3.1(+)-N-eGFP were bought from Genscript (NY, USA). The transfection was performed with jetPRIME® (0.8 µg DNA/dish) according to the manufacturer’s protocol. All cells were maintained in Dulbecco’s Modified Eagle’s Medium (DMEM) containing 10% fetal bovine serum. Twenty-four hours prior to the experiment, the cells were seeded at low density onto poly-D-lysine-coated dishes (35 mm Nunc Cell-culture dishes, Thermoscientific, USA)

### Membrane capacitance measurements using voltage square-wave stimulation

Recordings were performed in the whole-cell configuration using an Axopatch 200B amplifier and pClamp 10.2 software (MDS Analytical Technologies). A train of bipolar square-wave voltage pulses was applied with an amplitude ±40 mV and a frequency of 200 Hz. In the protocol the holding potential was set to 0 mV. Exponential current responses were low-pass filtered by a 10 kHz Bessel filter and digitized at 100 kHz using a Digidata 1440 (MDS Analytical Technologies). The cross-talk between R_S_ and C_M_ was suppressed by the following procedure: the acquired current traces were first deconvoluted with the transfer function of the recording apparatus. Passive membrane parameters of a cell were calculated from the theoretical function as described elsewhere (10). The pipette capacitance was recorded in the cell-attached mode first and subtracted from the currents recorded in the whole-cell configuration prior to further analysis. The patch pipettes used had a resistance of 2–4 MΩ. To stabilize the level of stray capacitance, the pipettes were coated with hydrophobic resin Sylgard184 (Dow Corning, USA). In the measurements the cells were continuously superfused with an external solution containing 140 mM NaCl, 3 mM KCl, 2.5 mM CaCl_2_, 2 mM MgCl_2_, 20 mM glucose, and 10 mM HEPES (pH adjusted to 7.4 with NaOH). For experiments with Cl^−^-free external solution, NaCl^−^ was replaced by Na^+^ MES^−^(methanesulfonate). The internal solution in the patch pipette contained 152 mM NaCl, 1 mM CaCl_2_, 0.7 mM MgCl_2_, 10 mM HEPES, and 10 mM EGTA (ethylenglycol-bis(aminoethylether)-N,N,N′,N′-tetra-acidic-acid) (pH 7.2 adjusted with NaOH). Ligands were applied via a four-tube or eight-tube ALA perfusion manifold using the Octaflow perfusion system (ALA Scientific Instruments, USA), which allowed for complete solution exchange around the cells within 100 ms. Currents were recorded at room temperature (20–24°C). A detailed description of the method can be found in (11).

### Membrane capacitance measurements with a dual-phase lock-in amplifier

Recordings were made using a dual-phase lock-in patch-clamp amplifier (SWAM-2C; Celica d.o.o., Ljubljana, Slovenia). Signals were low-pass filtered at 10 Hz with a 6-pole Bessel filter and digitized at 250 Hz by an A/D converter (CED 1401, Cambridge, UK) utilizing WCP software for acquisition (Dempster, University of Strathclyde, UK). HEK293 cells stably expressing hSERT were voltage-clamped at a holding potential of 0 mV, on which a sine-wave voltage (111 mV rms, 1600 Hz) was superimposed. In the compensated mode of recording, one of two outputs of the dual-phase lock-in amplifier signal is directly proportional to changes in C_M_. The correct phase angle was adjusted based on recordings elicited by the application of 1 pF calibration pulses. The patch pipettes used were coated with Sylgard184 and had a resistances of 2–4 MΩ. The employed solutions were the same as described above. For solution exchange, we used the Octoflow perfusion system (see above)

### Analysis of electrical circuits

Equivalent circuits were solved by loop analysis using Laplace transform. Below we show the analytical solutions for current differences before and after addition of a ligand in which inverse Laplace transform is indicated only formally. For the sake of space and clarity we used substitutions.

Inverse Laplace transform of a difference of two analytical solutions of the circuit shown in figure 2A and 6A is as follows:

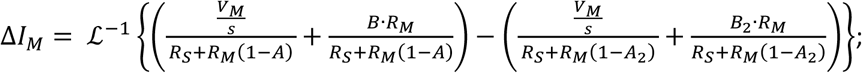

where:

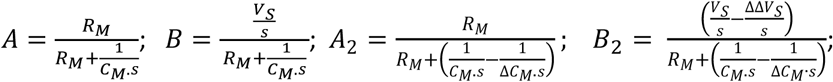

Inverse Laplace transform of a difference of two analytical solutions of the complex equivalent circuit shown on figure 6B is as follows:

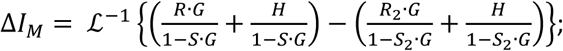

where:

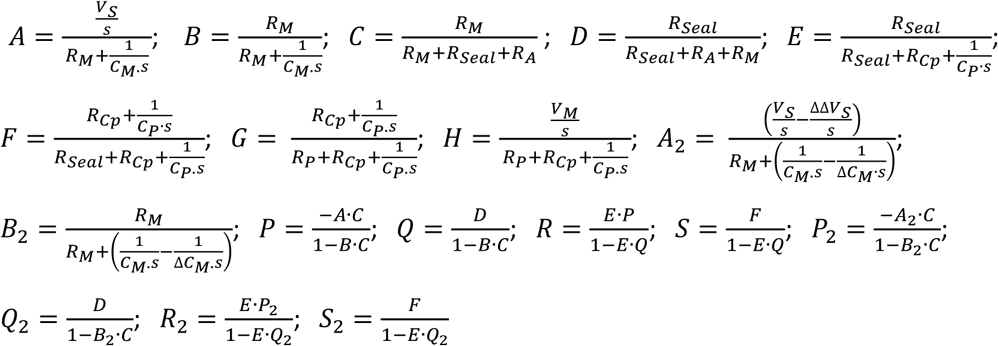

### Fitting procedure

Fitting of the residuals was performed with Matlab. We used the constrained version of the Matlab build-in function fminsearch (i.e. “fminsearchbnd.m” (D’Errico, 2012: https://de.mathworks.com/matlabcentral/fileexchange/8277-fminsearchbnd-fminsearchcon)) where the fitted parameter *ΔΔVS* and *ΔCM* were constrained to positive and negative values, respectively. The remaining parameters were set fixed and obtained from the individual recordings (C_M_, R_M_, R_A_, R_P_) and separate measurements (*Cp*= 3 pF). *R*_*Seal*_ was assumed to be 3 GΩ.

## Results

### Ligand induced capacitance changes are apparent changes

We previously showed that it is possible to monitor ligand binding to membrane proteins in real time by utilizing measurements of the cell membrane capacitance (C_M_)(8). We investigated the ligand-induced capacitance change in HEK293 cells stably expressing hSERT. Binding of a SERT ligand (e.g., cocaine) to the transporter resulted in a decrease in the total membrane capacitance (see Fig 1A -also shown in the figure is the time-dependent evolution of the circuit parameters R_M_ and R_S_). In Burtscher et al., we posited that this change in capacitance was caused by charged ligand binding to the protein. Upon adsorption to the binding site, the charge on the ligand becomes a surface charge (i.e. a fixed charge at the interface between the membrane and the surrounding aqueous solution). This offsets the electric potential at the outer surface of the membrane. As a consequence, the voltage difference between the inner and the outer membrane surface (ф_TRANS_) changes, which affects the ability of the membrane to store capacitive charges. Accordingly, the ligand-induced change in membrane capacitance (C_M_) does not result from a true capacitance change. It can rather be rationalized as follows. Approaches to measure the membrane capacitance rely on the assumption that the three-element circuit displayed in Fig. 1B can model the electrical response of a cell to a voltage stimulus (e.g. a square-wave voltage pulse - Fig. 1C&1D). However, one additional circuit component is needed in the equivalent circuit of the cell to represent changes induced by the adsorption of the charged ligand. This component is a battery, which must be added to account for changes of the charge densities at the inner and/or outer membrane surfaces. Omission of the battery from the circuit leads to an erroneous inference: the ligand-induced change in surface potential is misinterpreted as a change in the estimates of the other circuit components, most prominently of C_M_. This point is illustrated in Fig. 2. The figure shows an expanded circuit in which we incorporated an additional battery in series with the capacitor (Fig. 2A). The voltage (VS) at the battery (highlighted in red) was specified by the Gouy-Chapman model, which relates the potentials at the inner and outer membrane surfaces to the corresponding surface charge densities (Fig. 2B&C) (12, 13): Fig 2B shows the voltage profile over the membrane in the absence of cocaine at −40 mV and +40 mV, respectively. In this calculation, the initial inner and outer surface charge densities were assumed to be −0.018 C/m^2^ and −0.005 C/m^2^, respectively (14). The Gouy-Chapman model predicts that - due to the asymmetry of the inner and outer surface charge densities - VS increases by 6.4 mV (ΔVS), when the membrane potential (V_M_) is switched from −40 to +40 mV (i.e.by a voltage step of 80 mV). Importantly, the VS step is predicted to be oppositely directed to the V_M_ step (see Fig. 2A). Binding of cocaine to SERT adds one positive charge per occupied binding site and thus reduces the density of negative charges on the outer leaflet. Fig. 2C shows the voltage profile over the membrane in the presence of cocaine; again at −40 mV and +40 mV respectively. The calculation is based on the assumption that, on binding of cocaine, the outer surface charge density drops to −0.0034 C/m^2^. The difference of 0.0016 C/m^2^ is accounted for by the number of SERT moieties (i.e. 2*10^7^) expressed on the surface of our stable cell line (8), to which cocaine binds in a 1:1 stoichiometry at saturating concentrations. The Gouy-Chapman model predicts that ΔVS in the presence of cocaine is 7.9 mV, which differs by 1.5 mV from that in the absence of cocaine. The other circuit parameters (i.e. C_M_, R_M_ and R_S_) were chosen to closely approximate the circuit parameters of the experiment displayed in Fig 1A. Fig 2D shows the simulated current response of this circuit before (blue trace) and after (red trace) the application of a saturating concentration of cocaine. The evolution of the current over the first 10 µs was magnified in the inset to Fig 2D to show that the amplitude of the current response is slightly larger in the absence of cocaine. The simulated currents produced by the four circuit elements were then analyzed with the three-element circuit (Fig 2E). Because of this erroneous premise, this analysis inferred that all three circuit parameters had changed upon cocaine binding. While the predicted change in R_M_ was minute and not visible at the chosen scaling, the change in C_M_ and R_S_ were more pronounced. We want to emphasize, however, that the predicted change in R_S_ change fell within a range, where we observed spontaneous fluctuations in R_S_ in the majority of the actual measurements on cells. These variations in R_S_ were in most instances larger than the change predicted in the simulation. Moreover, in actual measurements we frequently observed an increase in R_M_ upon cocaine application (see also Fig. 1A). However, because we also observed a cocaine-induced R_M_ change in control cells (not shown) and because cocaine is known to inhibit ion channels, we attribute this small changes to a block of endogenous channels, which are present at low levels in our HEK293 cell line (15).

**Figure 1.**
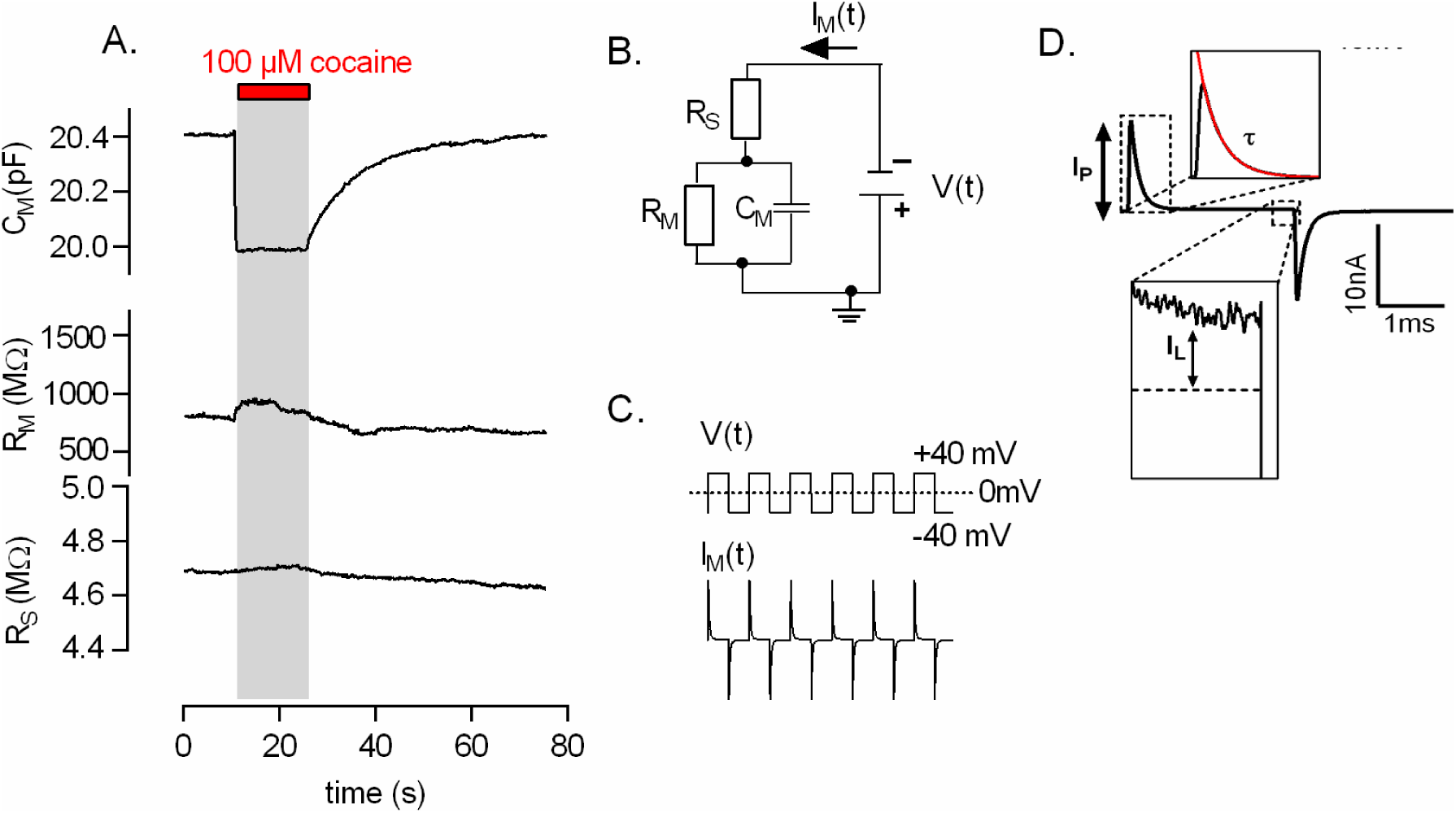
Cocaine binding to SERT reduces the total membrane capacitance (C_M_). **A.** A saturating concentration of cocaine (100 µM) was applied to a HEK293 cell stably expressing SERT and the change in membrane capacitance was recorded in the whole cell patch clamp configuration. Application of cocaine reduced the membrane capacitance. After 15 seconds the drug was removed from the bath solution, upon which C_M_ returned to initial values. Also shown is the time dependent evolution of the other circuit parameters (membrane resistance R_M_ and series resistance R_S_, which were obtained by fitting the current recordings elicited by the application of voltage square-wave pulses (**panel C**) to the equation describing the minimal equivalent circuit of the cell shown in (**panel B).** Data are from a representative experiment, which was reproduced 47 times. **D.** The circuit parameters are extracted by fitting a monoexponential function to the measured current decay. The fit (in red) yields three parameters (I_P_, τ and I_L_). I_P_ is the current amplitude at time point zero, τ is the time-constant of the current decay and I_L_ is the leak current following relaxation of the displacement current. With these parameters, it is possible to obtain R_S_ (R_S_= ΔV_M_/(I_P_+I_L_)), R_M_ (R_M_= ΔV_M_/I_L_-R_S_) and C_M_ (C_M_= τ*(1/R_S_+1/R_M_)).

### Ligand-induced capacitance changes assessed with sinusoidal voltage stimulation

In Burtscher et al. we measured the membrane capacitance with an approach, which relied on the analysis of current responses to voltage square pulses (8). More often, however, membrane capacitance measurements are conducted utilizing a sinusoidal voltage stimuli in combination with a hardwired (16) or a software-based lock-in amplifier (17). To see, if a change in the surface charge density of the membrane was also detectable with a lock-in amplifier, we simulated the current response of our extended equivalent circuit to a sinusoidal voltage stimulus (Fig.3A). In the simulation, we assumed that the cell was held at a holding potential of − 40 mV. At this holding potential, we superimposed a sinusoidal voltage stimulus with a frequency of 1 kHz and a peak-to-peak amplitude of 40 mV (see Fig. 3A and 3B). The Gouy-Chapman model was employed to calculate the voltage at the battery (ΔVS) in response to the sinusoidal change in V_M_, in time-steps of 1 microsecond (i.e. 1 MHz). The other circuit parameters were set to the values used in Fig. 2. Notably, the Gouy-Chapman model predicts that the voltage at the battery (ΔVS) is shifted by 90° with respect to the stimulating voltage (V_M_) (see Fig. 3A). Figure 3B shows the current response (I_M_) of this circuit to V_M_ (black trace) before (red trace) and after the application of cocaine (blue trace). As can be seen from the magnified traces in the inset to Fig. 3B, the simulation predicts a reduction in the amplitude of I_M_ upon cocaine binding. When subjected to admittance analysis, this change translated into a reduction in C_M_. Importantly, the calculated changes in the circuit parameters C_M_, R_M_ and R_S_ were exactly the same as predicted for the square-wave pulse method (Fig 2D). This suggests that both approaches to measure capacitance are ought to be sensitive to changes in surface potential. We experimentally verified the results of our simulation by recording cocaine-induced capacitance change from a cell expressing hSERT, by measuring the same with a dual-phase lock-in amplifier (see methods). As predicted, we were able to detect the cocaine-induced reduction in membrane capacitance. Fig.3C shows the two outputs of the amplifier. The upper panel displays the output that is directly proportional to changes in C_M_. It is evident that there is no cross-talk between the two channels. This shows that both approaches (square-wave pulse and sinusoid) are equally suited for monitoring ligand binding to membrane proteins.

**Figure 2.**
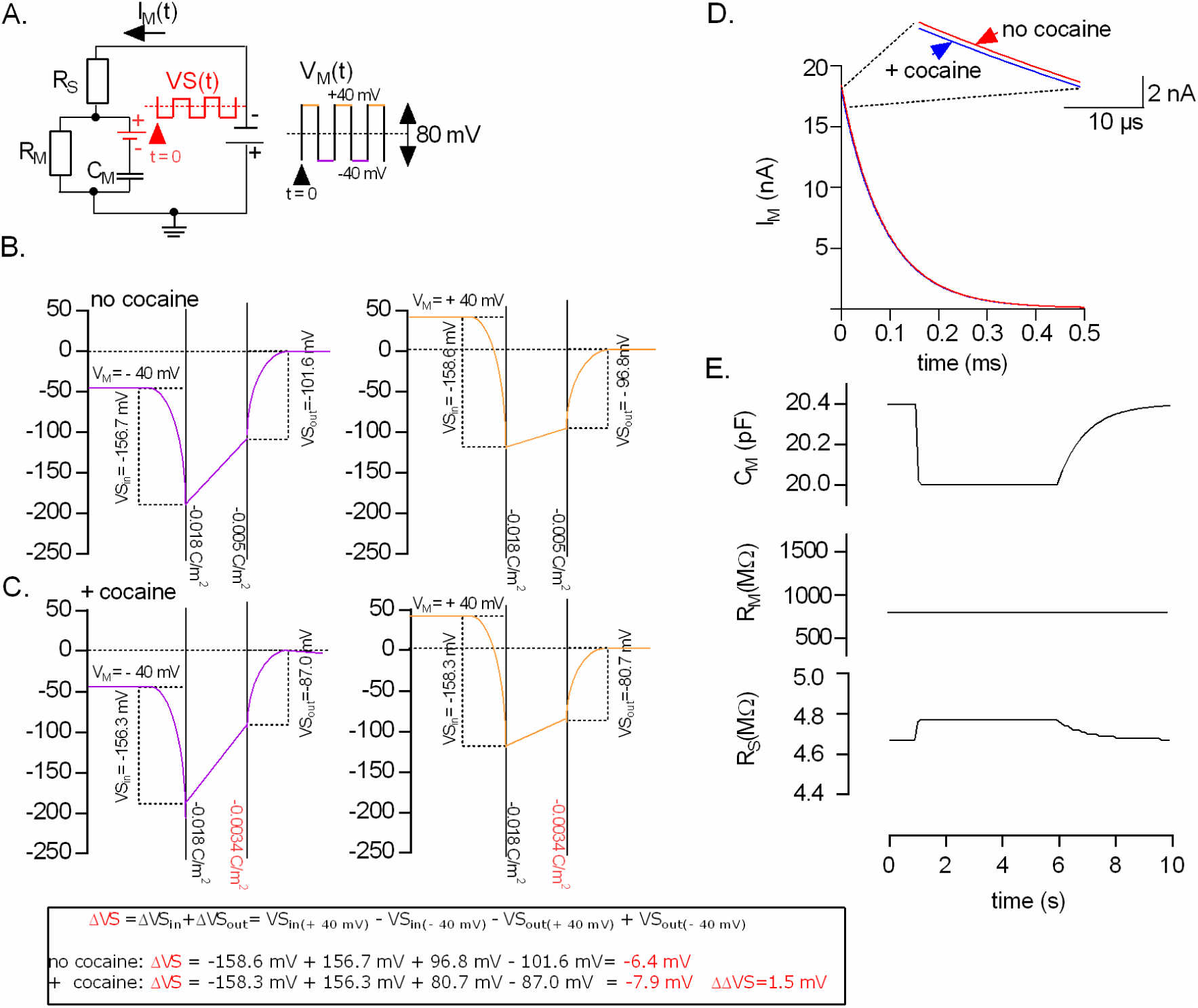
The apparent change in membrane capacitance can be accounted for by an extended equivalent circuit of the cell. **A.** Schematic representation of an extended equivalent circuit of the cell: a battery (in red) was added to the minimal equivalent circuit shown in Fig. 1B. The depicted voltage square-wave stimulus V_M_(t) was applied to the cell to elicit the recordings shown in Fig. 1A. V_M_ was stepped from − 40 mV to + 40 mV in time increments of 5 ms. The resulting changes in voltage VS(t) at the added battery were calculated by utilizing the Gouy-Chapman model and are shown as red traces. **B.** Voltage profile over the membrane in the absence of cocaine. The potential at the inner and outer surface were calculated utilizing the Gouy-Chapman model as adapted by (13). For the simulation we assumed that both the inner- and outer solution contained 150 mM NaCl. Initial surface charge densities for a HEK293 cell were taken from (14). The left and right hand panel in B show the voltage profile for voltage differences (V_M_) between the outer and inner bulk solution of − 40 mV and +40 mV, respectively. **C.** Voltage profile over the membrane in the assumed presence of a saturating concentration of cocaine. The left and the right panel show the voltage profile at − 40 mV and + 40 mV, respectively. The potential at the membrane surfaces in C are different from that in B. This is because in C the outer surface charge density was assumed to change to − 0.0034 C/m^2^ as a consequence of cocaine binding. The inserted box at the bottom of panel C shows for the calculation of the voltage step, which occur at the added battery according to the Gouy-Chapman model. **D.** Simulated current responses of a cell expressing SERT in the assumed absence (red trace) and presence (blue trace) of cocaine. The inset shows that the current amplitude of the response in the absence of cocaine is slightly larger than in the presence of cocaine. **E.** Simulated current responses were analyzed by employing the minimal equivalent circuit of the cell (Fig. 1B). This analysis misinterprets the change at the battery as a change in the other circuit parameters. The parameters which are most affected are R_S_ and C_M_.

**Figure 3.**
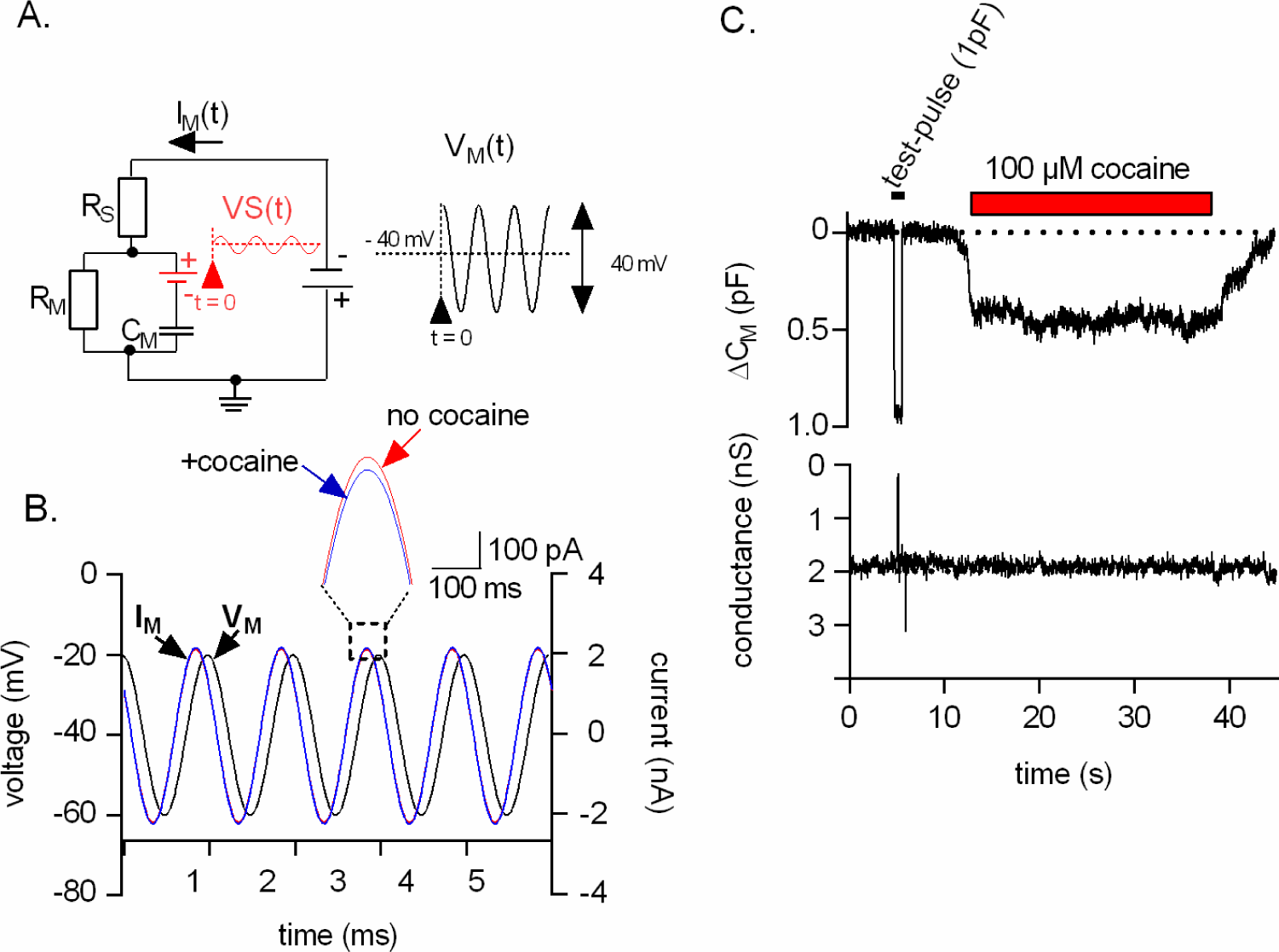
Effect of a sinusoidal voltage stimulus on the extended equivalent circuit. **A.** The voltage change at the battery VS(t) (red trace) was calculated using a simulation. This simulation assumed that a sinusoidal voltage V_M_ (t) (black trace) with a peak to peak amplitude of 40 mV and superimposed on a holding potential of −40 mV was applied to the extended equivalent circuit. The frequency of the applied sinusoid was 1 kHz. The voltage at the battery VS(t)) was specified by the Gouy-Chapman model, which predicted a shift in VS by 90° wiht respect to V_M_. **B.** The current response (I_M_) to the stimulating voltage (V_M_-black trace) was extracted from simulations assuming the absence (red trace) and the presence (blue trace) of cocaine. The inset shows a magnified section of the currents (I_M_): the amplitude of the current in the absence is larger than in the presence of cocaine. **C.** The change in membrane capacitance C_M_ was recorded in the whole cell patch clamp configuration from a HEK293 cell stably expressing hSERT using a hard wired lock-in amplifier. Application of 100 µM cocaine (indicated by the red bar) led to a reduction of C_M_. Upon cocaine washout, C_M_ relaxed to initial values.

### The simulated current response to a voltage square pulse differs depending on whether the change is a true or an apparent capacitance change

A change in any of the circuit parameters defining the equivalent circuit of the cell must result in a shape change of the current response to a square-wave voltage pulse. However, the characteristics of this shape change depend on the underlying alteration in the circuitry. This point is illustrated in Fig. 4: the left hand and the middle panel of Fig 4A show the simulated current response to a square-wave voltage pulse in the absence and presence of cocaine, respectively. In this simulation, cocaine binding was accounted for by a 1.5 mV change in the voltage (ΔΔVS) supplied by the battery. The right hand panel shows the subtraction of the two traces which for the following description will be referred to as the “residual”. The residual corresponds to the cocaine-induced change in current, which we denote as the “battery residual”. In Fig. 4B, we convoluted the battery residual with the impulse response function of our recording apparatus to emulate the residual of an actual measurement. The impulse response function was acquired as described in (10, 11). In Fig. 4C, we show the overlay of the battery residual with the simulated residual of a true capacitance change (referred to as “capacitive residual” henceforth), which was obtained by changing C_M_ in the simulation (i.e. ΔC_M_ =1pF). It is evident from the overlay of the two residuals that they differ in their kinetics. For instance, the rising phase of the “battery residual” is faster than the rising phase of the “capacitive residual “. The reason for this is as follows: changes of the voltage supplied by the battery give rise to residuals that are only subject of the intrinsic filters of the recordings apparatus and not of the filter constituted by the cell (i.e. tau∼R_S_*C_M_, cutoff-frequency=1/tau). In contrast, in patch clamp experiments the current required to charge the membrane (i.e. the displacement current) must pass through R_S_, which results in a slow rise of the “capacitive residual”.

**Figure 4.**
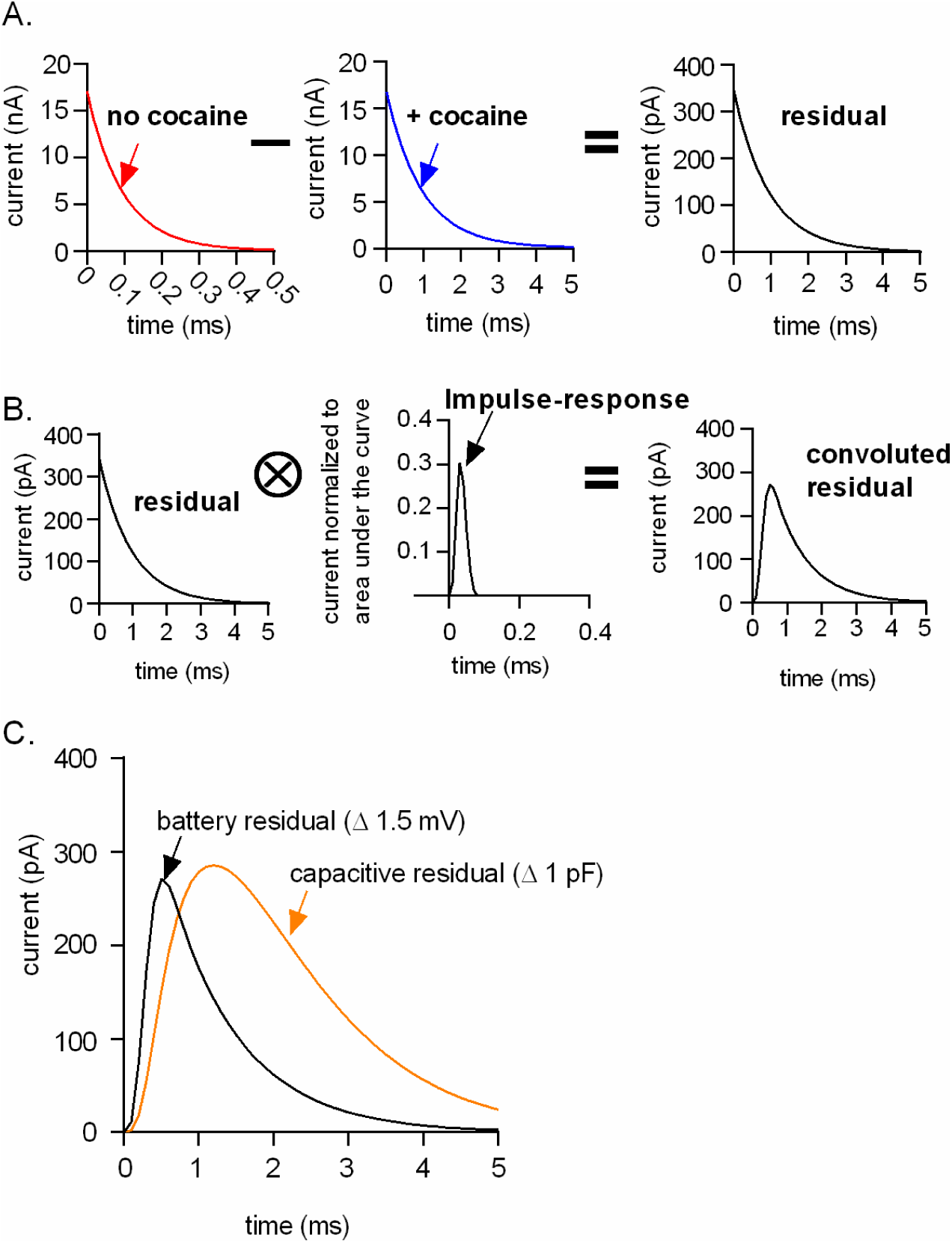
The shape of the differential current (i.e. residuals) produced by square-wave voltage pulses depends on the underlying alteration in the circuitry. **A.** The left hand and the middle panel show the simulated current response of the extended circuit depicted in Figs. 2A and 3A in the absence (red trace) and presence of cocaine (blue trace), respectively. The right hand panel shows the subtraction of the two traces, i.e. the residual. **B.** Convolution of the simulated residual (left) with the impulse response function obtained from the recording apparatus employed (middle panel). The right hand panel shows the convoluted residual produced by the change at the battery. **C.** Comparison of the residual produced by the change at the battery - i.e. the battery residual - with the simulated residual of a true capacitance change (orange trace).

### Model residuals measured utilizing the compensation circuitry of the patch clamp amplifier

Fig. 5A shows an overlay of two residuals measured with our recording apparatus; one is a residual of a true capacitance change and, the other, of an apparent capacitance change. We generated the residual of the true capacitance change by using the compensation circuitry of the patch clamp amplifier (Axopatch 200B). We emulated the current response of a typical HEK293 cell by setting the whole cell capacitance and series resistance compensation to 26 pF and 8 MΩ, respectively. Currents were elicited by the application of voltage steps (V_M_ =80 mV). The capacitive residual was obtained by changing the compensation setting to 25.5 pF and subtracting the resulting current from a current recorded at the prior setting (i.e. 26 pF). The battery residual was recorded as follows: we kept the compensation circuitry settings at 26 pF and 8 MΩ and recorded the current in response to an 80 mV step pulse. We then changed the stimulus voltage by 1 mV (i.e. 79 mV) to mimic the ligand induced change in ф_TRANS_. The displayed residual (red trace) was obtained by subtracting the two current responses (i.e. V_M, 80 mV_ –V_M, 79 mV_). It is evident that the measured residuals match the simulated residuals shown in Fig. 4C. We analyzed the currents produced by the compensation circuitry utilizing the three-element circuit. As expected, the modeled true capacitance change was correctly identified as a sole change in C_M_ (Fig. 5B). However, the analysis showed that the error made by assuming an unchanged V_M_ (i.e. 80 mV instead of 79 mV) had translated into a change in R_S_ and C_M_ (Fig. 5C). R_M_ in both measurements was infinite.

**Figure 5.**
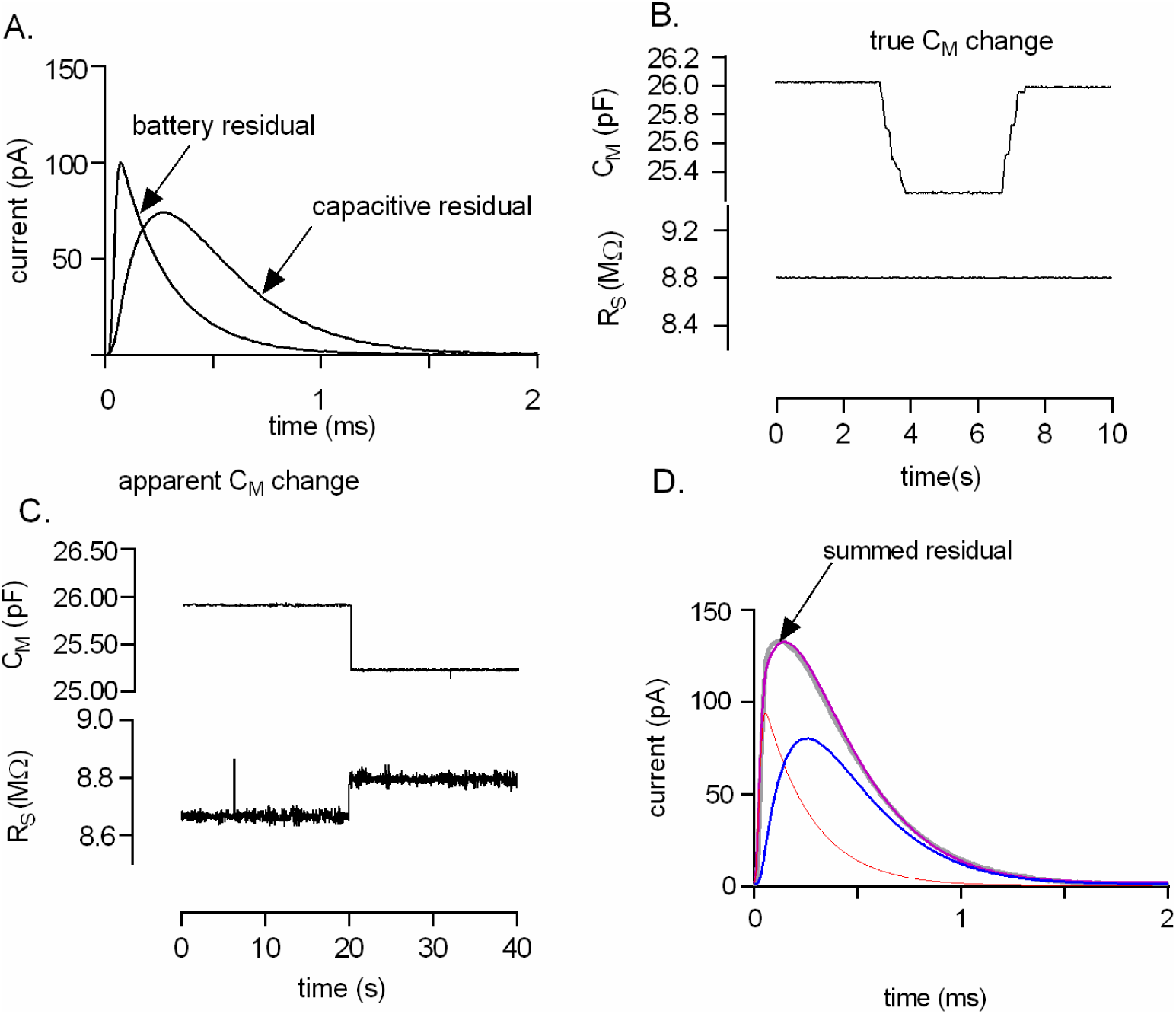
Measured residuals utilizing the compensation circuitry of the patch clamp amplifier. A. Modeled battery and capacitive residual (see text for a detailed description of the approach employed to obtain the parameters). **B.** Analysis of the recorded true capacitance change using the minimal equivalent circuit of the cell. **C.** Analysis of the modeled apparent capacitance change using the minimal equivalent circuit of the cell. **D.** The summed measured residuals of A (grey trace) were fitted to an analytical solution of the extended circuit (trace in magenta). The fit allowed for decomposing the summed residual into a battery (orange trace) and capacitive residual (blue trace).

### Analysis of mixed responses

So far, we assumed that a measured capacitance change can either originate from a true change in capacitance or from an apparent change (i.e., from alterations in surface potential). However, a scenario can be envisaged, in which a manipulation (e.g. application of a ligand or a voltage change) triggers a mixed response. We reasoned that - because of the distinct shape of the two types of residuals - each individual component can be isolated even in those instances, where they occur simultaneously. We tested our ability to decompose a mixed response by summing the measured residual displayed in Fig 5A and subjected the sum to the following fitting procedure: an analytical expression describing the response of the extended equivalent circuit to a square pulse was derived from the Laplace transformation (18). The sum of the residuals was fitted by using the difference of two analytical solutions. To take into account the filters in our signal path, we transformed this difference into the frequency domain using fast Fourier transformation and convolved it with the impulse response of the recording apparatus. The filtered difference of the analytical solutions was then back-transferred into the time domain using inverse fast Fourier transform and was fitted to the experimental data with only two fitting parameters: (i) Δ C_M_ which describes a difference in C_M_ and (ii) ΔΔVS, which accounts for the change in surface potential. The other circuit parameters (C_M_, R_S_, R_M_), were fixed and the corresponding values were estimated by fitting the current responses to the three-element circuit. As can be seen from Fig. 5D, the fit (shown in magenta) was able to decompose the summed residual (grey trace) into its original elements, i.e. the battery residual (in red) and the capacitive residual (in blue). Thus, this approach provides the means for monitoring both processes and their evolution over time in instances where true capacitance changes and changes in surface potential occur in parallel.

### Analysis of the current responses recorded from mammalian cells

In the upper panel of Fig. 6A we show a residual (grey trace) recorded from a HEK293 cell stably expressing SERT, which originated from cocaine binding. The noise level in the residual acquired from the cell was higher than in the residuals produced by the compensation circuitry (cf. Fig. 5A). This was presumably due to the higher impedance noise and due to spontaneous fluctuations in the circuit parameters, both of which seem inherent to recordings conducted on cells. There is an additional difference between the residual produced by the compensation circuitry and the recorded data from the cell: in contrast to the former, the cellular residual was recorded utilizing a glass electrode, which acted as a capacitor, because it was immersed into the bath solution. This introduced an additional low-pass filter into the signal path. As a result, the circuit shown in the lower panel of Fig.6A (i.e. simple model) failed to adequately account for the time course of the rising phase of the residual. This is evident from the inset in Fig 6A in which we show the first 100 µs of the recording: in this time interval the fit (green trace) deviates considerably from the data.

**Figure 6.**
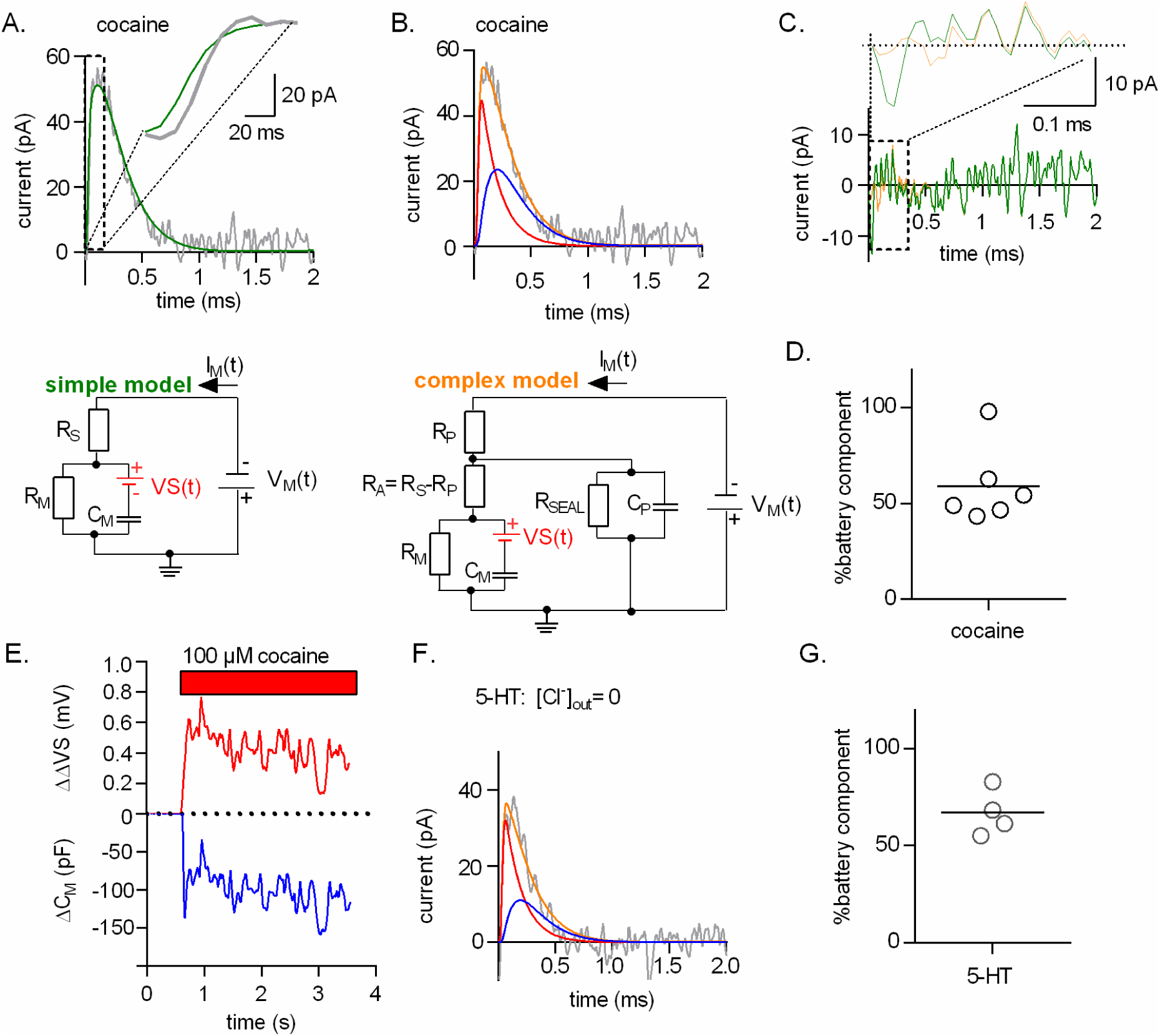
Residuals measured from HEK293 cells stably expressing SERT cells challenged with cocaine or 5-HT can be decomposed into a battery and a capacitive residual. **A.** The upper panel shows a residual obtained from recordings of a HEK293 cell stably expressing SERT, which was superfused with cocaine binding (grey trace). The measured residual was fitted to the simple circuit shown in the lower panel. The inset shows the magnified rising phase of the residual. The fit (in green) did not adequately account for the initial phase of the residual. The lower panel depicts the simple model **B.** The upper panel shows the same residual as in panel A fitted to the analytical solution of a more complex circuit displayed in the lower panel of B. The fit (in orange) allowed for decomposing the measured residual into a battery (in red) and a capacitive residual (in blue). The lower panel depicts the complex model **C.** Comparison of the fitted residuals of the simple model (in green) and the complex model (in orange). The inset shows the magnified initial phase of the fit. In the initial phase the more complex model better accounted for the data. **D.** Contribution of the battery component in % to the total change in C_M_ induced by cocaine. Shown is the summery of 6 independent experiments. In average the contribution of the battery component was 60 % ± 20 %. **E.** Change in ΔΔVS over time as a result of cocaine binding plotted along with the simultaneous change in C_M_. **F.** Measured residual of a HEK293 cell stably expressing SERT, which was challenged with 30 µM serotonin (5-HT) in the absence of extracellular Cl^−^(in grey). The residual was fitted to the complex model. The fit (orange trace) decomposed the measured residual into a battery (in red) and a capacitive residual (in blue). **G.** Summary of 4 independent experiments in which 5-HT was applied. Plotted in % is the contribution of the battery residual to the total change in capacitance (67.0 % ± 11.9 %).

In protocols utilizing voltage square-wave pulses, the frequency richness allows for modeling the cellular current response by more complex equivalent circuits (19). In the circuit shown in the lower panel of Fig. 6B (i.e. complex model), we introduced additional circuit elements that influence cellular recordings. These are the stray capacitance of the electrode (C_P_), its resistance (R_P_) and the seal resistance (R_SEAL_). We derived an analytical solution of this circuit (see methods) using Laplace transform. We fitted the residual, which had resulted from the cocaine-induced change by the same procedure as outlined before, except that we employed the model shown in the lower panel of Fig. 6B instead of the model in the lower panel of Fig.6A. We emphasize, however, that - regardless of the model employed - we only allowed two parameters to vary (i.e. Δ C_M_ and ΔΔVS). The other parameters were fixed in both models. In the case of the complex model, we obtained the required circuit parameter values as follows: initial values for C_M_, R_M_ and R_S_ were extracted by analyzing the current response in the absence of cocaine with the three-component circuit shown in Fig. 1B. R_P_ was measured prior to every experiment by the application of a square-wave voltage pulse to an electrode immersed into the bath solution. For estimating C_P_, we recorded the current responses to square wave voltage pulses from 3 different Sylgard-coated pipettes immersed into vaseline to emulate a gigaOhm-seal. C_P_-values were calculated by integrating the current responses to obtain the charge (Q) and by division of Q with the applied voltage (C_P_ =3pF). R_A_ is the difference between R_S_ and R_P_. R_Seal_ was assumed to be 3 GΩ. In Fig. 6B, we show the fit of the complex model (orange trace) to the data. In Fig. 6C, we plotted the fitted residuals of the simple model (in green) and the complex model (in orange) for comparison. Both models were able to adequately account for the larger part of the recorded residual, however the complex model was superior in fitting the rising phase of the residual (see inset in Fig. 6C).

The fit revealed that the residual, which had originated from cocaine binding to SERT is comprised of a battery (red trace) and a capacitive component (blue trace) (Figure 6B-upper panel). This was unexpected and disproved our original claim that the cocaine-induced change in C_M_ was fully attributable to changes in surface potential (8). However, in partial exoneration of this claim, we found that the battery residual accounted in average for a slightly larger part of the change. In Fig. 6D, we show the analysis of 6 independent experiments. This figure indicates the relative contribution of the apparent capacitance change of each experiment to the total capacitance change. In average we found that about 60% of the change had been caused by a ligand-induced alteration in the outer surface charge density. The remaining 40% represented a true capacitance change. It is currently unclear why cocaine binding also gave rise to a true capacitance change. We surmise that cocaine binding to SERT may have induced conformational rearrangements, which reduced the polarizability of the protein. In Fig. 6E, we show the time-dependent evolution of the cocaine-induced true capacitance change (ΔC_M_) and of the transmembrane potential change (ΔΔVS). This figure demonstrates that our analysis allows for individually tracking both changes when they occur simultaneously. In Fig. 6F, we show another example of a residual recorded from a cell stably expressing hSERT. In this experiment, we applied 30 µM of the endogenous substrate 5-HT in the absence of extracellular Cl^−^. Extracellular Cl^−^ was omitted because this prevented substrate induced conformational rearrangements while still allowing for binding of the substrate to SERT (8). The fit (in orange) decomposed the residual that had originated from binding of 5-HT to SERT into a battery (in red) and a capacitive component (in blue). Fig. 6G summarizes the analysis of 4 independent experiments showing that the C_M_ change induced by 30 µM 5-HT is to the larger part (65%) caused by adsorption of the charged ligand to SERT.

It is implicit to our model that ligand-induced changes in surface potential occur as a consequence of charged ligand binding. In contrast, the binding of an uncharged ligand is expected to be ineffective in altering the outer surface potential and consequently in producing a “battery residual”. This prediction is difficult to verify utilizing HEK293 cells expressing SERT, because all ligands known to bind to this transporter harbor a positive charge. However, other SLC-transporters exist, which bind neutral (i.e. zwitter-ionic) ligands. Notable examples are the transporters for glycine (i.e. GlyT1 and GlyT2). Accordingly, in the present study we employed Cos7 cells transiently expressing GlyT1 and GlyT2, respectively. We used Cos7 cells, because unlike HEK-293 cells, they did not express endogenous glycine transporters. We then asked the question how binding of an uncharged ligand (i.e. glycine) to glycine transporters impinges on the measured membrane capacitance. Accordingly, we recorded the capacitance of cells expressing GlyT1 and GlyT2, which were challenged by 1 mM glycine. Fig. 7A and 7B show the result of typical measurements obtained from cells expressing GlyT1 and GlyT2: application of glycine led to a reduction in membrane capacitance in cells expressing either of the two glycine transporter subtypes. Upon subsequent removal of glycine from the bath solution the capacitance relaxed to initial values. Plotted in these figures are also the evolution of R_M_ and R_S_ over time. When tested in control cells, application of glycine did not result in an appreciable change in the membrane capacitance (data not shown)

**Figure 7.**
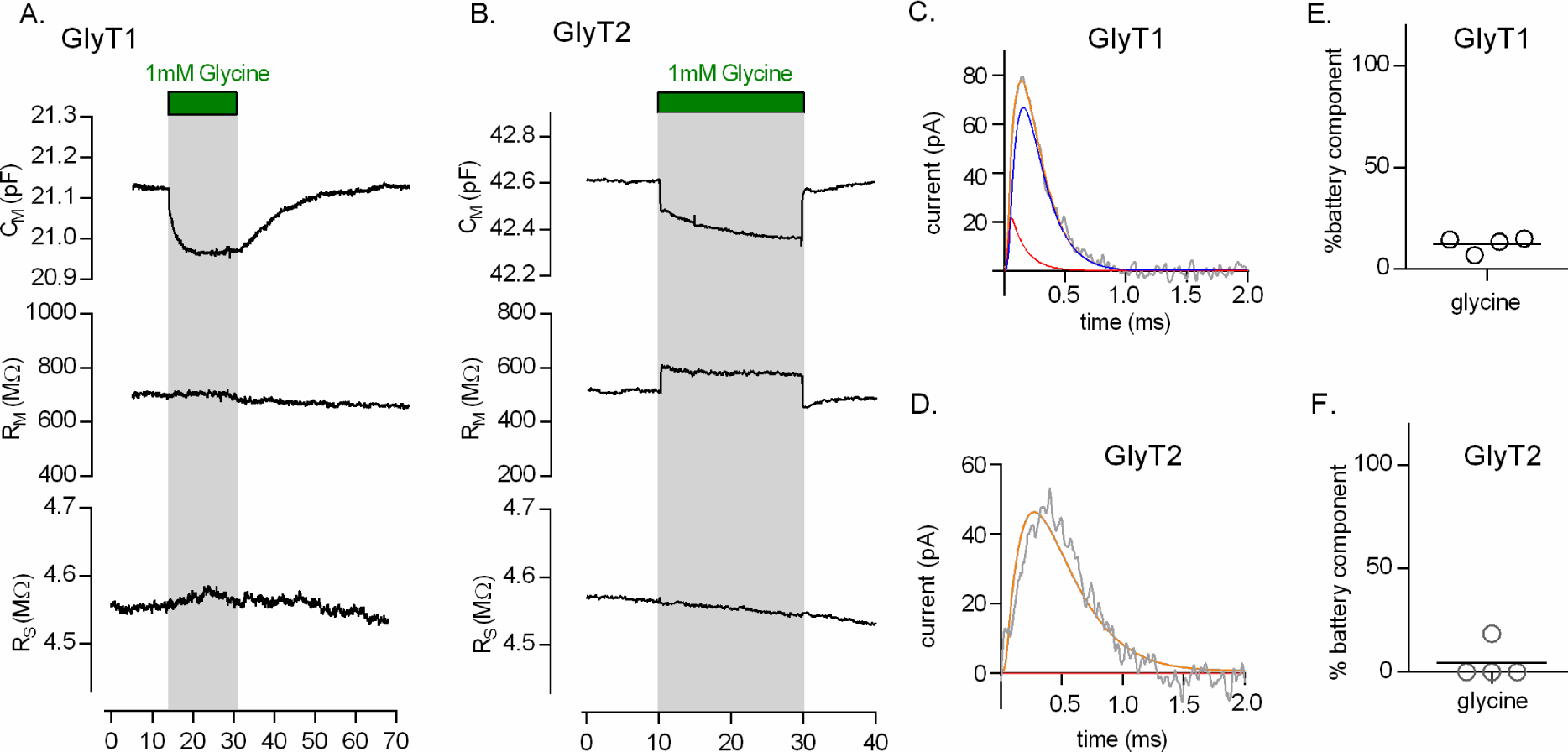
Application of glycine reduces the membrane capacitance of COS-7 cells expressing GlyT1 and GlyT2. **A & B.** The change in membrane capacitance C_M_ was recorded in the whole cell patch clamp configuration a COS-7 cell transiently expressing GlyT1 (A) or GlyT2 (B), which were challenged by the application of 1 mM glycine. The time-dependent evolution of the circuit parameters is shown: in both GlyT1 (A) and GlyT2 expressing cells, application of 1mM glycine gave rise to a reduction in C_M_. However R_S_ and R_M_ were not affected by glycine in panel A but glycine increased R_M_ in panel B. After 15 seconds, glycine was removed from the bath solution. Upon removal of the amino acid, C_M_ and R_M_ relaxed to initial values. **C.** Measured residual produced by the application of glycine to a COS-7 cell expressing GlyT1. The fit (orange trace) of the data (grey trace) to the complex model shown in Fig. 6B decomposed the residual into a small battery (in red) and into a large capacitive residual (in blue). **D.** Measured residual of a COS-7 cell expressing GlyT2 produced by binding of 1 mM glycine (grey trace). The complex model only inadequately accounted for the data (orange trace). The fitted parameter - ΔΔVS - converged to zero. Accordingly, there is no battery residual. **E. and F.** show in % the contribution of the battery residual to the total change in capacitance for GlyT1 (12.4 % ± 3.8 %) and GlyT2 (4.6 % ± 9.3 %), respectively (n=4).

Inspection of the residual produced by the application of 1 mM glycine to cells expressing GlyT1 revealed that the underlying change had mostly been caused by a true change in C_M_ (see Fig.7C). The contribution of the battery residual amounted to only about 12%, suggesting that glycine binding had only minimally affected the outer surface potential (see Fig. 7E). This result was in line with the net zero charge of glycine in the physiological pH-range.

In cells expressing GlyT2 the observations were similar (see Fig 7D & 7F). However, the residuals derived from GlyT2 expressing cells challenged with glycine were even slower than predicted for a true change in C_M_ (see Fig.7D). It was reported that GlyT2 but not GlyT1 bound extracellular Na^+^ in a voltage dependent manner. Voltage jumps from negative (i.e. − 60 mV) to more positive potentials resulted in outwardly directed transient currents, which were presumably carried by dissociating Na^+^ ions (20). Our data suggest that the voltage at the membrane settles faster than Na^+^ can unbind. However, the model does not consider Na^+^ binding. We surmise that it is for this reason why the fit failed to adequately account for the data. We note that the application of glycine let to an increase in R_M_ (Fig.7B). This effect of glycine was absent in control cells (data not shown). We therefore also attribute this change to slow Na^+^ dissociation from GlyT2. Slow dissociation of Na^+^ is expected to affect the fit-parameter-I_L_ (see Fig.1D) and thereby R_M_. Synchronized Na^+^ unbinding from GlyT2 is prevented by glycine binding, which we believe is the reason why R_M_ is increased in the presence of the amino-acid (20).

## Discussion

We previously showed that it is possible to detect changes of the surface potential at the plasma membrane by measurements of the membrane capacitance (8). In the present study, we demonstrate that, in addition, it is also possible to distinguish between a change in surface potential (i.e. apparent capacitance change) and a true capacitance change. Capacitance measurements rely on the assumption that the electric properties of a recorded cell are adequately described by a three-element electric circuit. However, this minimal equivalent circuit of the cell cannot account for a change in the cellular surface potential. In order to model the latter it is necessary to use a more complex circuit that contains a battery connected in series with the capacitor. This complex circuitry predicts that the shape of the differential current (i.e. residual) produced by a true and an apparent capacitance change, respectively differ. We experimentally verified this prediction and we confirmed that the cocaine and 5-HT induced change measured in cells expressing SERT were indeed and to a larger part caused by alterations in the surface potential. We also showed that the electrically neutral ligand glycine only minimally affected the surface potential of the plasma membrane when bound to glycine transporter 1 or glycine transporter 2 (i.e. GlyT1 and GlyT2). These findings are consistent with the concept that - upon binding to a membrane protein - the charge of the ligand becomes a surface charge and thereby offsets the outer surface potential.

The electrical potentials at the inner and the outer surface of the plasma membrane are to a large extent defined by fixed charges at the interface between the membrane and the surrounding aqueous solution (21). These arise primarily from the net negative charge of phospholipid head groups but also from water-accessible acidic or basic amino acid residues within membrane proteins. In addition, charged molecules, which bind to the plasma membrane or to membrane proteins also contribute to the surface charge. Although both membrane surfaces carry a negative net charge, the net negative charge on the inner surface normally exceeds that on the outer surface (22). This is mainly due to the asymmetric distribution of membrane lipids between the inner and the outer membrane leaflet (e.g. the inner membrane leaflet shows a higher content of phosphatidylserine). This asymmetry is maintained by specialized proteins, which are tasked with moving phospholipids from the inner to the outer membrane layer (i.e. flippases) or in the opposite direction (i.e. floppases) (23, 24). These proteins harness the energy required to maintain the lipid/charge asymmetry from the hydrolysis of ATP (25). The energy stored in this lipid gradient can be readily dissipated by scramblases, which are proteins that facilitate diffusive lipid crossing between the two membrane leaflets (26, 27).

Surface charges play an important role in many biological processes. For instance, because of their net-negative surface charge, cells tend to repel each other. This zeta potential prevents cell aggregation (28). In fact, the zeta potential is the basis for the single file mode of capillary transit in vivo and the slow sedimentation of red blood cells ex vivo: the erythrocyte sedimentation rate has been used for more than a century as a sensitive - albeit non-specific - parameter to detect inflammation and other abnormalities in people. The net-negative surface charge of cells is also one of the reasons, why specific interactions between cell surface proteins are required for establishing cell contacts. The adhesive forces provided by these proteins are needed to overcome the electrostatic repulsion. Moreover surface charges greatly affect the concentrations of charged solutes in the vicinity of the membrane. Within a perimeter of about 1 nm, the concentrations of positively charged or negatively charged molecules are much higher or lower, respectively, than in the bulk solution (29). In addition, the asymmetric distribution of surface charges (i.e. excess of negative charge on the inner surface) increases the driving force for cellular uptake of positively charged solutes but reduces the driving force for uptake of negatively charged solutes. Both effects are expected to shape the intra-and extracellular concentration of signaling molecules such as monoamines or glutamate. In addition, surface charges are known to impinge on the voltage dependence of voltage-gated ion channels (30). By virtue of this action, they gauge the excitability of cells such as cardiomyocytes, skeletal muscle cells and neurons (31).

It is evident from this non-exhaustive list of examples that many cellular functions are affected by changes of the inner and/or the outer surface charge densities. However, the mechanisms, which regulate the surface charge densities at the plasma membrane, are poorly understood. This is in part due to lack of experimental approaches that allow for detecting changes in surface charges in real time. Here, we show that it is possible to track alterations in the surface potential by an electrophysiological approach. Importantly, the method described in the present study allows for discriminating between a change in surface potential and a change in true capacitance, even if these occur in parallel. This can be of relevance if a process is monitored that gives rise to simultaneous changes in the surface area of the plasma membrane and the surface charge density. This can plausibly be the case during mitosis, apoptosis, T cell activation etc. (32–34). Likewise, changes in the surface charge density may also occur in parallel with changes that affect the polarizability of a membrane protein. In this context, it is worth noting that cocaine binding to SERT also resulted in a true capacitance change. Specifically, the capacitance was reduced when cocaine was bound to SERT. This implies that the polarizability of the cocaine bound state of SERT is lower than that of the apo-state. It is unclear, however, how the macroscopically observed change can be explained on the microscopic level. At the current stage, the underlying conformational changes in SERT remain elusive and difficult to address by an experimental approach. However, we anticipate that this will be amenable to computational studies in the near future because of the continuous progress in structural studies of SERT.

## Author Contributions

V. B., M. H, M.F and W.S designed the experiments and wrote the paper. V. B. performed the experiments shown in Fig.1, Fig.3, Fig.5, Fig.6 and Fig.7.. M.H and W.S. performed the simulations in Fig.2, Fig.3, Fig.4 and Fig.6.. M.H. scripted the fitting routine used in Fig.5, Fig.6 and Fig.7. All authors reviewed the results and approved the final version of the manuscript.

## Acknowledgments

We thank Marjan Slakrupnik for his help with the dual-phase lock-in amplifier. We thank Shreyas Bhat for proofreading this manuscript and Klaus Schicker for helpful discussions. This work was supported by the Austrian Science Fund (FWF) grant P28090 and P31813 to WS and the WWTF-grant LS17-026 to MF.

The authors declare no competing financial interests.

## References

1. Lindau, M., and E. Neher. 1988. Patch-clamp techniques for time-resolved capacitance measurements in single cells. Pflügers Arch. Eur. J. Physiol. 411: 137–146.

2. Von Gersdorff, H., and G. Mathews. 1994. Dynamics of synaptic vesicle fusion and membrane retrieval in synaptic terminals. Nature. 367: 735–9.

3. Fernandez, J.M., E. Neher, and B.D. Gomperts. 1984. Capacitance measurements reveal stepwise fusion events in degranulating mast cells. Nature. 312: 453–455.

4. Bean, B.P., and E. Rios. 1989. Nonlinear Charge Movement in Mammalian Cardiac Ventricular Cells. Components from Na and Ca Channel Gating. J. Gen. Physiol. 94: 65–93.

5. Kilic, G., and M. Lindau. 2001. Voltage-dependent membrane capacitance in rat pituitary nerve terminals due to gating currents. Biophys. J. 80: 1220–9.

6. Lu, C.C., a Kabakov, V.S. Markin, S. Mager, G. a Frazier, and D.W. Hilgemann. 1995. Membrane transport mechanisms probed by capacitance measurements with megahertz voltage clamp. Proc. Natl. Acad. Sci. U. S. A. 92: 11220–11224.

7. Gilson, M.K., and B.H. Honig. 1986. The dielectric constant of a folded protein. Biopolymers. 25: 2097–119.

8. Burtscher, V., M. Hotka, Y. Li, M. Freissmuth, and W. Sandtner. 2018. A label-free approach to detect ligand binding to cell surface proteins in real time. Elife. 7: e34944.

9. Kevin D. Gillis. 1995. Techniques for Membrane Capacitance Measurements. In: Single-Channel Recording.

10. Hotka, M., and I. Zahradnik. 2017. Reconstruction of membrane current by deconvolution and its application to membrane capacitance measurements in cardiac myocytes. PLoS One. 12: e0188452.

11. Burtscher, V., M. Hotka, and W. Sandtner. 2019. Detection of Ligand-binding to Membrane Proteins by Capacitance Measurements. BIO-PROTOCOL. 9: pii: e3138.

12. Genet, S., R. Costalat, and J. Burger. 2000. A few comments on electrostatic interactions in cell physiology. Acta Biotheor. 48: 27.

13. Plaksin, M., E. Shapira, E. Kimmel, and S. Shoham. 2018. Thermal Transients Excite Neurons through Universal Intramembrane Mechanoelectrical Effects. Phys. Rev. X. 8: 011043.

14. Zhang, P.C., A.M. Keleshian, and F. Sachs. 2001. Voltage-induced membrane movement. Nature. 413: 428–32.

15. Wright, S.N., S.Y. Wang, Y.F. Xiao, and G.K. Wang. 1999. State-dependent cocaine block of sodium channel isoforms, chimeras, and channels coexpressed with the β1 subunit. Biophys. J. 76: 233–45.

16. Neher, E., and A. Marty. 1982. Discrete changes of cell membrane capacitance observed under conditions of enhanced secretion in bovine adrenal chromaffin cells. Proc. Natl. Acad. Sci. 79: 6712–6716.

17. Gillis, K.D. 2000. Admittance-based measurement of membrane capacitance using the EPC-9 patch-clamp amplifier. Pflugers Arch. Eur. J. Physiol. 439: 655–64.

18. Widder, D. 1941. The Laplace Transform. Princeton, NJ: Princeton University Press.

19. Thompson, R.E., M. Lindau, and W.W. Webb. 2001. Robust, high-resolution, whole cell patch-clamp capacitance measurements using square wave stimulation. Biophys. J. 81: 937–48.

20. López-Corcuera, B., R. Martínez-Maza, E. Núñez, M. Roux, S. Supplisson, and C. Aragón. 1998. Differential properties of two stably expressed brain-specific glycine transporters. J. Neurochem. 71: 2211–9.

21. McLaughlin, S.G.A. 2004. Divalent Ions and the Surface Potential of Charged Phospholipid Membranes. J. Gen. Physiol. 58: 667–87.

22. Zachowski, A. 2015. Phospholipids in animal eukaryotic membranes: transverse asymmetry and movement. Biochem. J. 294: 1–14.

23. Groen, A., M.R. Romero, C. Kunne, S.J. Hoosdally, P.H. Dixon, C. Wooding, C. Williamson, J. Seppen, K. Van Den Oever, K.S. Mok, C.C. Paulusma, K.J. Linton, and R.P.J. Oude Elferink. 2011. Complementary functions of the flippase ATP8B1 and the floppase ABCB4 in maintaining canalicular membrane integrity. Gastroenterology. 141: 1927–37.

24. Contreras, F.X., L. Sánchez-Magraner, A. Alonso, and F.M. Goñi. 2010. Transbilayer (flip-flop) lipid motion and lipid scrambling in membranes. FEBS Lett. 584: 1779–86.

25. Zachowski, A., J.P. Henry, and P.F. Devaux. 1989. Control of transmembrane lipid asymmetry in chromaffin granules by an ATP-dependent protein. Nature. 340: 75–6.

26. Sahu, S.K., S.N. Gummadi, N. Manoj, and G.K. Aradhyam. 2007. Phospholipid scramblases: An overview. Arch. Biochem. Biophys. 462: 103–14.

27. Kodigepalli, K.M., K. Bowers, A. Sharp, and M. Nanjundan. 2015. Roles and regulation of phospholipid scramblases. FEBS Lett. 589: 3–14.

28. Fernandes, H.P., C.L. Cesar, and M. de L. Barjas-Castro. 2011. Electrical properties of the red blood cell membrane and immunohematological investigation. Rev. Bras. Hematol. Hemoter. 33: 297–301.

29. Green, W.N., and O.S. Andersen. 1991. Surface Charges and ION Channel Function. Annu. Rev. Physiol. 53: 341–59.

30. Madeja, M. 2000. Extracellular Surface Charges in Voltage-Gated Ion Channels. News Physiol Sci. 15: 15–19.

31. Han, P., B.J. Trinidad, and J. Shi. 2015. Hypocalcemia-induced seizure: Demystifying the calcium paradox. ASN Neuro. 7: pii: 1759091415578050.

32. Ma, Y., K. Poole, J. Goyette, and K. Gaus. 2017. Introducing membrane charge and membrane potential to T cell signaling. Front. Immunol. 8: 1513.

33. Adam, G., and G. Adam. 1975. Cell surface charge and regulation of cell division of 3T3 cells and transformed derivatives. Exp. Cell Res. 93: 71–8.

34. Taruvai Kalyana Kumar, R., S. Liu, J.D. Minna, and S. Prasad. 2016. Monitoring drug induced apoptosis and treatment sensitivity in non-small cell lung carcinoma using dielectrophoresis. Biochim. Biophys. Acta - Gen. Subj. 1860: 1877–83.

